# LTR transposable elements contribution to the apple genome, methylome and transcriptome evolution in *Malus domestica*

**DOI:** 10.64898/2026.09.17.750989

**Authors:** Andréa Bouanich, Gwendoline Couturier, Nathalie Choisne, Maryline Cournol, Angélina El Ghaziri, Charles-Elie Rabier, Jean-Marc Celton, Claudine Landès

## Abstract

The whole genome duplication (WGD) shared by apple (*Malus domestica*) and pear (*Pyrus communis*) dated 27 Mya was followed 21 Mya by a burst of transposable elements (TE), making these organisms a prime choice for studying the evolution of duplicated genes. In this study, we conducted a meta-analysis of 149 RNA-Seq datasets and focused on gene pairs for which one ohnolog was systematically under-expressed compared to its duplicate. To understand this systematic differential expression, we investigated the TE environment (TE type, TE divergence and insertion position) of these genes pairs and found that under-expressed genes in apple and pear were enriched in recent Class I LTR TE Copia and Gypsy insertions in their immediate genic environment. In apple, we identified a quantitative relationship between the number of LTR insertions in the genes’ environments and the level of differential expression among pairs of ohnologs. Finally, we found that these under-expressed genes displayed either hypomethylated or hypermethylated profiles in their sequence in the CG and CHG contexts. Altogether, our results highlight the major role played by TE in shaping genome structural evolution, and their contribution to gene transcription regulation, with potential consequences on species divergence and emergence of new phenotypic traits.

## 1 Introduction

Whole genome duplication (WGD) is a common phenomenon in the evolution of eukaryotic organisms in both animals (Lynch and Connery 2000; J. Zhang 2003; Jaillon et al. 2004) and plants (Wendel 2000; Soltis et al. 2004; Van de Peer 2011; Panchy et al. 2016). While the ancestor of all seed plants underwent a WGD about 319 Mya, a second WGD occurred in the ancestor of all flowering plants about 192 Mya (Cui et al. 2006; Jiao et al. 2011). Duplicated genes resulting from WGD are called ohnologous genes, in honor of Susumu Ohno, who predicted that WGDs played a major role in vertebrate evolution (Ohno 1970). He hypothesized that duplicated genes may evolve via neofunctionalization, subfunctionalization, or conserve their original function.

Today, we know that most duplicated genes undergo a process known as pseudogenization or loss through chromosomal rearrangements (Zhe Yu et al. 2020). Therefore, determining the evolutionary forces responsible for the maintenance of duplicated genes is key to understanding the evolutionary adaptation and emergence of novel traits. Numerous population genetics models have been developed to address this question.

We refer to Kusmin et al. 2022 for a recent review of the mechanisms underlying the conservation of duplicated genes. Moreover, the most recent models and bioinformatics tools are described in the reviews by Hahn 2009 and Lallemand et al. 2020, respectively.

Following WGD, an increase in transposable elements (TE) content known as TE burst (Vicient and Casacuberta 2017) has been observed in many organisms such as *Capsella bursa-pastoris* (Agren et al. 2016), *Thinopyrum intermedium* (Divashuk et al. 2016) and *Brassica* (An et al. 2014). This burst results in numerous random insertions of TEs in the genome, impacting duplicated genes evolution via gene loss, neofunctionalization, or chromosomal rearrangements as observed in maize (Bruggmann et al. 2006). Among TEs, Class I Long Terminal Repeat (LTR) Retrotransposons (copy-and-paste replication mechanism) can range from a few hundred bases to over 5 kb. They have two associated superfamilies, Copia and Gypsy, which differ in the succession of their RT (Reverse Transcriptase), RH (RNase H) and INT (Integrase) domains (Wicker et al. 2007).

The *Rosaceae* family displays a high diversity of flowers and fruits, in part due to the numerous WGDs in this clade (Xiang et al. 2017; Soundararajan et al. 2019). The most recent known WGD in the *Maleae* clade is shared by apple, pear, rowan, and quince (H. Li et al. 2019).

Apple, pear, and rowan trees are native to the Tian Shan mountains in the Himalayan mountain chain (formed from 25 to 30 Mya) (S. A. Harris et al. 2002; Silva et al. 2014; M. Li et al. 2017), which results from the collision of the Asian and Indian plates starting approximately 55 Mya (Vance and N. Harris 1999; Aitchison et al. 2007). The elevation of the mountains caused a significant climatic change for the local species, with variations in temperature, UV exposure, and atmospheric pressure (Valdiya 1999). These climatic changes could have triggered the last WGD (Van de Peer et al. 2021) dated 27 Mya (Lallemand et al. 2023).

Following an autopolyploid WGD (Hodel et al. 2022), chromosomal rearrangements and a return to a diploid state, *Maleae* now display a 2n=34 genome (Daccord et al. 2017) as compared to the 2n=18 genome of the ancestor (Illa et al. 2011). The Golden Delicious Double Haploid #13 (GDDH13 v1.1) apple genome and the W65 Double Haploid pear genome (Linsmith et al. 2019), both display 17 chromosomes, with 50,522 and 37,445 annotated genes, respectively. While the pear genome has not been annotated for repeated sequences, the apple genome contains up to 59.5% of repeated sequences (Lallemand 2022), mostly TE.

According to the classification of Wicker et al. 2007, most TE in the apple genome correspond to Class I TE (42.9%) while Class II TE represent a smaller proportion of the genome (13.4%). Analysis of TE sequences revealed that this genome has undergone two TE bursts, estimated at 21 Mya and at 1.7 Mya (Daccord et al. 2017). In many plants that have undergone allopolyploid WGD, subgenome dominance has often been observed (e.g. *Arabidopsis thaliana* Freeling and Thomas 2006, *Zea mays* Schnable et al. 2011, *Brassica rapa* X. Wang et al. 2011).

In a previous study, Lallemand et al. 2023 showed that TE coverage varied among pairs of GDDH13 ohnologous chromosomes, and identified a potential subgenome dominance in this autopolyploid species. This dominance was identified via a significant imbalance in mapped QTLs between ohnologous chromosomes. However, no difference was observed in the rate of evolution between the coding sequences of ohnologous genes. An imbalance in the number of differentially expressed gene pairs between ohnologous chromosomes was observed on the basis of 149 RNAseq experiments derived from several apple cultivars and species, tissues and experiments.

In this study, we investigated the impact of post-WGD TE insertions on the differential expression of ohnologous genes, and how distinct TE environments impacted the methylation profiles associated with these genes in apple. To confirm our findings, we used pear as a biological replicate of apple. We also studied whether the accumulation of Class I and LTR order TE insertions correlated with the differential expression levels observed among pairs of ohnologous genes in apple. Finally in apple, we investigated the methylome of groups of ohnologous genes and characterized particular patterns of methylation, some of them associated with specific TE families insertion.

## 2 Methods

### 2.1 W65 pear genome TE annotation

As performed for the apple genome (Daccord et al. 2017) and to facilitate the comparison between apple and pear, the TEdenovo pipeline from the REPET package v2.5 (Quesneville et al. 2003; Edgar and Myers 2005; Quesneville et al. 2005; Flutre et al. 2011; Amselem et al. 2019) was used on the pear W65 genome to detect TEs and to provide a consensus sequence for each TE family. Consensus TE sequences were then used to annotate the TE copies in the whole genome using the TEannot pipeline38 from the REPET package v2.5. Manual curation of the results was then performed in order to validate potentially uncertain annotations.

### 2.2 Differential expression analysis among pairs of ohnologous genes in pear

First, the identification of syntenic genes within ohnologous chromosomes was performed using the i-ADHoRe 3.0 (Proost et al. 2012) ad hoc pipeline as described in Lallemand et al. 2023, allowing the reconstruction of syntenic fragments displayed using Circos (Krzywinski et al. 2009). Secondly, differential transcription level among pairs of ohnologous genes was estimated as described in Lallemand et al. 2023. This differential expression analysis was performed using 53 publicly available pear RNA-seq experiments each containing three biological replicates, and three RNA-seq experiments each containing two biological replicates. To complete this design, we generated RNA-seq data from five additional tissues harvested from the W65 genotype, including leaves, young leaves, buds, stems and flowers (ENA:PRJEB112848).

### 2.3 Characterization of differential expression gene groups in apple and pear

#### Differential expression gene groups definition

As previously described in Lallemand et al. 2023, using data derived from apple and pear RNA-seq experiments (149 and 61 for apple and pear, respectively), we counted the number of occurrences a gene was differentially over-expressed relative to its ohnolog. Based on these counts we categorized groups of ohnologous gene pairs according to their differential expression. Pairs in which one gene was systematically over-expressed compared to its ohnolog in all RNA-seq experiments were referred as non switching gene pairs. Pairs in which the over-expressed gene varied within the pair in at least 19 percent of the experiments in apple, and in at least 28 percent of the experiments in pear, were referred as switching pairs. We selected 19 and 28 percent as thresholds because they yielded numbers of ohnologous pairs similar to the numbers of non-switching pairs in apple and pear, respectively. Pairs in which both ohnologs were never expressed in all RNA-seq experiments were called Not Expressed (NE). This analysis focuses only on duplicated pairs of ohnologs from the last WGD.

#### Gene ontology enrichment analysis

Gene Ontology (GO) (Ashburner et al. 2000) enrichment analyses were performed for two sub-ontologies: Biological Process (BP) and Molecular Function (MF).

To achieve this enrichment, we first updated the functional annotation of the GDDH13 v1.1 and W65 genes by homology to *Arabidopsis thaliana*. A local all by all protein BLAST (blastp) (Altschul et al. 1990) was performed between the 2 proteomes using the following parameters: e-value at 0.001 with a search for 5 homologs and default matrix BLOSUM62. The best BLAST hit was retained. GO terms of *Arabidopsis thaliana* retrieved from the TAIR11 database (Cheng et al. 2017) downloaded in April 2025 were then added. To perform GO terms enrichment analyses, the non switching gene group for each species was divided into two groups: over-expressed and under-expressed. GO term enrichments of gene groups (non switching, switching, and NE) were then conducted using the QuaDS pipeline (Bouanich et al. 2025). QuaDS allows the characterization of qualitative variables (group of individuals, cluster, etc.) from qualitative and/or quantitative variables. This tool automatically analyzes the relationship between each descriptive variable and the target variable. For this analysis, the significance level was set at 0.01 for the various statistical tests (Chi-square test of independence, Fisher’s exact test, factor levels description according an hypergeometric distribution, and v-test statistic). The categorical variables to be analyzed were the GO sub-ontologies (BP, MF, CC) and GO terms, and the variables to be characterized were our gene groups (non switching, switching, and NE).

### 2.4 Transposable Element analysis

For both apple and pear genomes, the three detection algorithms, PILER (Edgar and Myers 2005), GROUPER (Quesneville et al. 2003), and RECON (Bao and Eddy 2002), were used to identify repeated sequences. We also considered these algorithms to estimate TE divergence. TE sequences aligned with these consensus sequences were tagged either as (a) very recent TEs (PILER algorithm, 99%-100% identity), (b) recent TEs (GROUPER algorithm, 95%-99% identity), or (c) old TEs (RECON algorithm, 50%-95% identity).

A new TE annotation of the double haploid pear W65 genome was performed. The distributions of TE identity percentages were analyzed according to their order. The relative frequency of each identity percentage was calculated by dividing the number of occurrences by the total number of elements belonging to the corresponding order. The resulting distributions were then represented as frequency curves as a function of identity percentage. The pear bursts datations were estimated based on the TE bursts in apple published in Daccord et al. 2017 and on Kimura’s divergence (“International Human Genome Sequencing Consortium” 2001).

We then investigated the TE environment within 5kb of the genes using the TEGRIP pipeline (Meguerditchian et al. 2021), which provides information on the relationship between the gene and the TE.

Using start and end of genes’ and TEs’ positions, the pipeline returns 6 possible gene/TE relationships (TE upstream, TE downstream, overlapping upstream or downstream, TE in the gene sequence, and TE containing the gene) as represented in Figure 1.

**Fig. 1:**
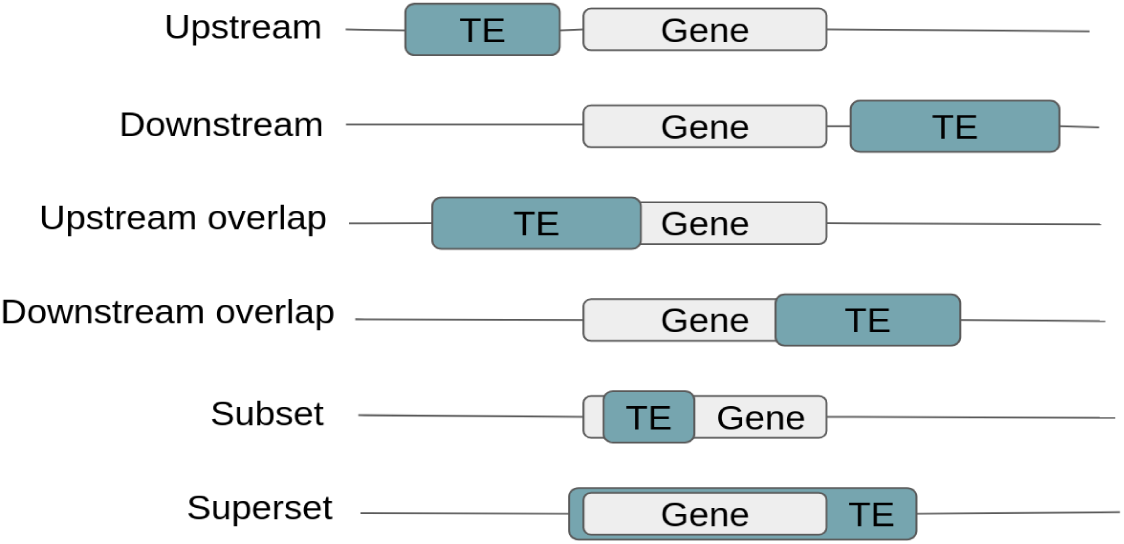
Representative scheme of the different gene/TE relationships that can be obtained using the TEGRIP pipeline.

We then investigated whether differentially expressed gene groups were enriched in particular TE environments. To this end, we separated the non switching genes into two groups, the non switching over-expressed and the non switching underexpressed. Using the QuaDS pipeline (Bouanich et al. 2025), analyses were then performed to investigate potential over- or under-representations of TE environments by comparing the 4 different gene groups (non switching over-expressed, non switching under-expressed, switching, and NE). Variables such as TE Class, Order, and Superfamily were analyzed. Enrichment analyses were also conducted using data from consensus building algorithms (very recent, recent, and old). The last descriptive variable taken into account in the enrichment is the gene/TE relationship. The following statistical tests were performed thanks to the QuaDS pipeline: Chi-square test of independence, Fisher’s exact test, factor levels description (according to an hypergeometric distribution) with 0.05 significance level.

Finally, in apple, we investigated a potential effect of the number of LTR insertions (Gypsy or Copia) on the differential expression levels of ohnologs within the non switching group, using a linear mixed model. For this analysis, we subdivided the non switching gene pairs into nine categories. These categories ranged from category 1, representing no LTR insertions in either of the two non switching genes’ environments, to categories 8 and 9, representing two or more LTR insertions in one of the two non switching genes’ environments. Details of the categories are in Supplementary data STab1. The mixed model (Boisgontier and Cheval 2016) was applied to expression data from non switching gene pairs derived from the 149 RNA-seq experiments investigated. The mixed model parameters were as follow: LTR category was treated as fixed effect, gene pairs was treated as the random effect, and LogFC as the dependent variable. The p-values were adjusted for multiple testing with the FDR BH procedure (Benjamini and Hochberg 1995).

### 2.5 Methylation profiles analysis in apple

We used the Bioconductor package BPRmeth method (Binomial Probit Regression for Methylation) (Kapourani and Sanguinetti 2016; Kapourani and Sanguinetti 2018) with the original implementation in version 1.28 to quantify the level of methylation on specific genomic regions using BS-seq data derived from leaf samples from GDDH13 (Daccord et al. 2017). At a given location, the methylation intensity level corresponds to the proportion of reads containing a cytosine over the total number of reads (see Kapourani and Sanguinetti 2016). The BPRMeth method is a kernel-based statistical learning method. This method is used to estimate methylation profiles, which correspond to the methylation intensity curve along a nucleic acid sequence. This distribution may vary according to the methylation context and the genomic regions analyzed.

We analyzed separately the methylation level in the contexts (a) CG, (b) CHG and (c) CHH. Furthermore, for each methylation context, we considered two genomic regions: (1) from 500 bases upstream to the beginning of the gene, (2) in the gene sequence (containing introns). Regions with less than 5 cytosines were discarded. Then, BPRmeth clusters genes with similar methylation profiles into groups. The optimal number optimal groups was estimated using the Bayesian Information Criterion (BIC) (Neath and Cavanaugh 2012).

Next, the methylation profiles were tagged either hypermethylated, hypomethylated, or variable. Profiles were assigned as hypermethylated or hypomethylated if at least 75% of the genomic region was above 0.75 of below 0.25 methylation level, respectively. Other profiles were tagged as variable.

We then investigated whether particular methylation profiles were over or under-represented in the four gene groups (non switching over-expressed, non switching under-expressed, switching, and NE). Using the QuaDS pipeline, an enrichment analysis was performed with the same parameters as for the TE enrichment analysis. For genes lacking a minimum of 5 cytosines, we assigned arbitrarily a hypomethylated profile. To further the research, an analysis of gene groups methylation was conducted, taking into account the TE environment of these genes. To this end, the gene groups were divided into subgroups based on the TE present in their environment. Finally using QuaDS we investigated whether the insertion of a TE class (Class I, Class II or unknown class) or order (LTR, LINE, TIR or unknown order) next to gene subgroups correlated with particular methylation profiles.

## 3 Results

### 3.1 Pear genome TE annotation

The 3,292 TE consensus sequences provided by the TEdenovo detection pipeline were used to annotate their copies in the whole genome. In the W65 genome, TEs represent 52.9% of the genome sequence for a total of 318,639 TEs. The most abundant repeats in this genome are retrotransposons of Class I elements (38.2%), followed by Class II elements (8.9%), and TE of unknown Class (5.2%). The abundance of repeat sequences by order is described in Supplementary data SFig1.

### 3.2 Pear ohnologous genes identification

Using i-ADHoRe, we identified a total of 13,415 pairs of ohnologous genes in the pear genome distributed along 454 syntenic blocks. Of these, based on the analysis of the highest number of contiguous syntenic blocs along chromosomes, reflecting the most recent WGD, we identified a total of 8,403 pairs of ohnologous genes derived from the last WGD (Supplementary data SFig2).

### 3.3 Differential expression gene groups

Among the 16,779 ohnologous pairs found in apple by Lallemand et al. 2023, we retained the 10,819 pairs from the last WGD belonging to the ohnologous chromosome pairs: 01-07; 01-15; 02-07; 02-15; 03-11; 04-06; 04-12; 05-10; 06-14; 08-15; 09-17; 12-14; and 13-16. The remaining 5,960 pairs of genes are distributed in other smaller syntenic fragments that likely originate from older WGD.

In apple, the analysis derived from the RNA-seq data allowed us to identify 828 non switching gene pairs, 837 switching gene pairs, and 74 NE genes pairs. In pear, we identified 203 non switching gene pairs, 189 switching gene pairs, and 20 NE pairs. In both apple and pear, the non switching group was further divided into two groups of genes, *i.e.* the non switching over-expressed group and the non switching under-expressed group (Table 1).

**Table 1:**
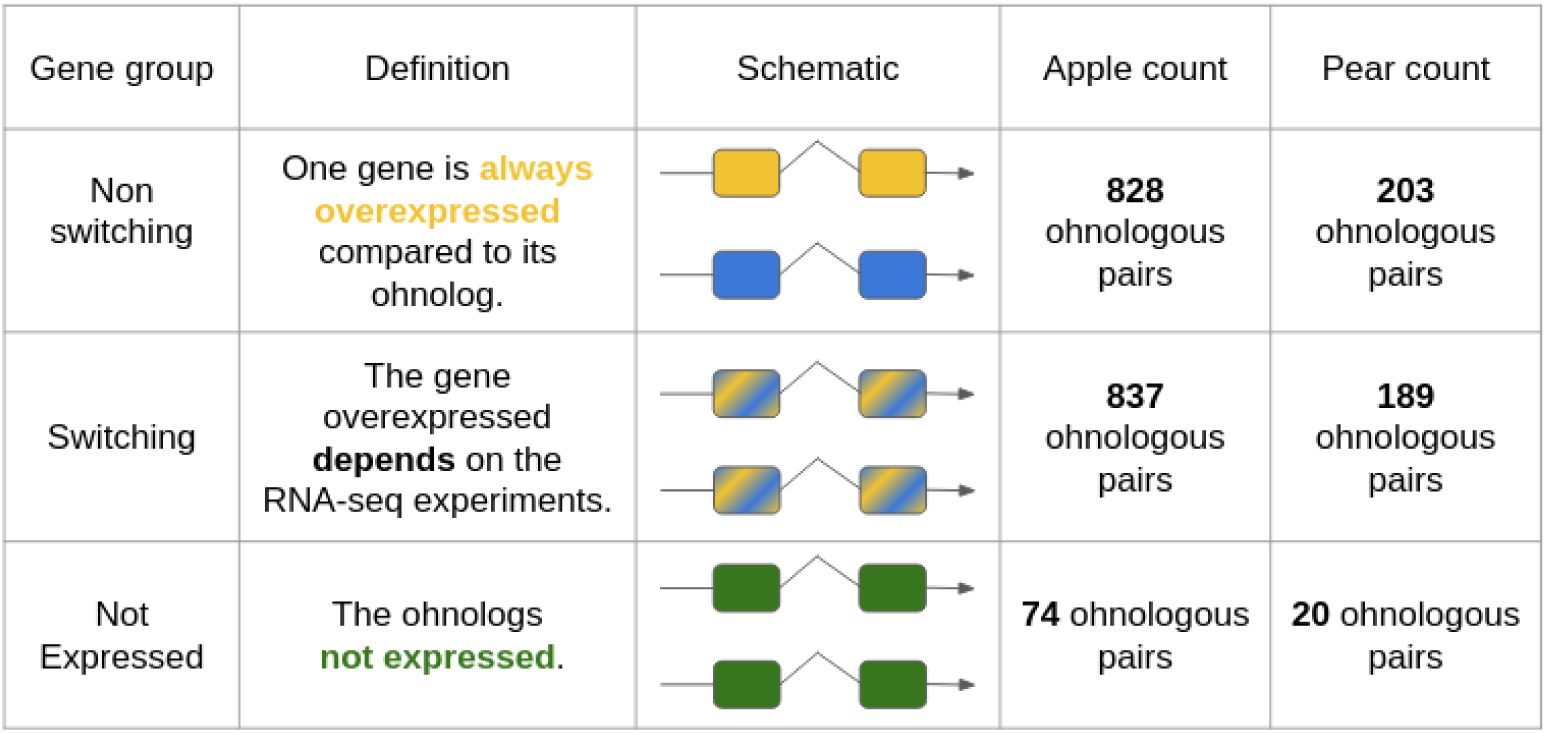
Description of the different gene groups: Non-switching genes are represented either in yellow (always over-expressed) or blue (always under-expressed) color, in the 149 and 61 RNAseq experiments in apple and pear, respectively. Switching genes, which differential expression vary between ohnologues are represented in both yellow and blue colors, and Not expressed genes are represented in green color

| Gene group | Definition | Schematic | Apple count | Pear count |
| --- | --- | --- | --- | --- |
| Non switching | One gene is **always overexpressed** compared to its ohnolog. |  | **828** ohnologous pairs | **203** ohnologous pairs |
| Switching | The gene overexpressed **depends** on the RNA-seq experiments. |  | **837** ohnologous pairs | **189** ohnologous pairs |
| Not Expressed | The ohnologs **not expressed**. |  | **74** ohnologous pairs | **20** ohnologous pairs |

### 3.4 GO term enrichments

For apple, enrichment analyses indicated that non switching genes were enriched in Molecular Function GO terms associated with gene expression regulation, at the translational and post-transcriptional levels, involved in chromatin dynamics and ribosome biogenesis. Non switching genes were also enriched in Biological Process GO terms associated with cell regulation, organelle function, and embryonic development. NE genes were enriched in Molecular Function GO terms associated with plant development and morphogenesis, as well as transcriptional regulation, cell signaling, stress responses, and hormone signaling. Several Biological Process GO terms indicate the involvement of these genes in specialized biosynthetic pathways. These genes are also involved in post-translational modification and protein interactions. Switching genes were found to be enriched in Molecular Function GO terms associated with transcriptional regulation and activation of gene expression. They are involved in Biological Process GO terms associated with protein phosphorylation, cell signalling, cell development and differentiation (Supplementary data STab2).

For pear, enrichment analyses indicated that non switching genes were enriched in Molecular function GO terms associated with mRNA binding, while switching genes were enriched in GO terms associated with transcription factor activity. NE genes were enriched in Molecular function GO terms associated with enzymatic activities and signalisation (Supplementary data STab2).

### 3.5 Transposable Elements analysis

TE bursts in apple have already been described in (Daccord et al. 2017). In this study, we investigated a potential TE burst in pear. The identity percentages of TE copies were represented at the TE order level in Figure 2. Two bursts can be observed at 80% and at 94% identity. The first and largest burst was estimated at 18 Mya and the second one, smaller, was estimated at 2.7 Mya.

**Fig. 2:**
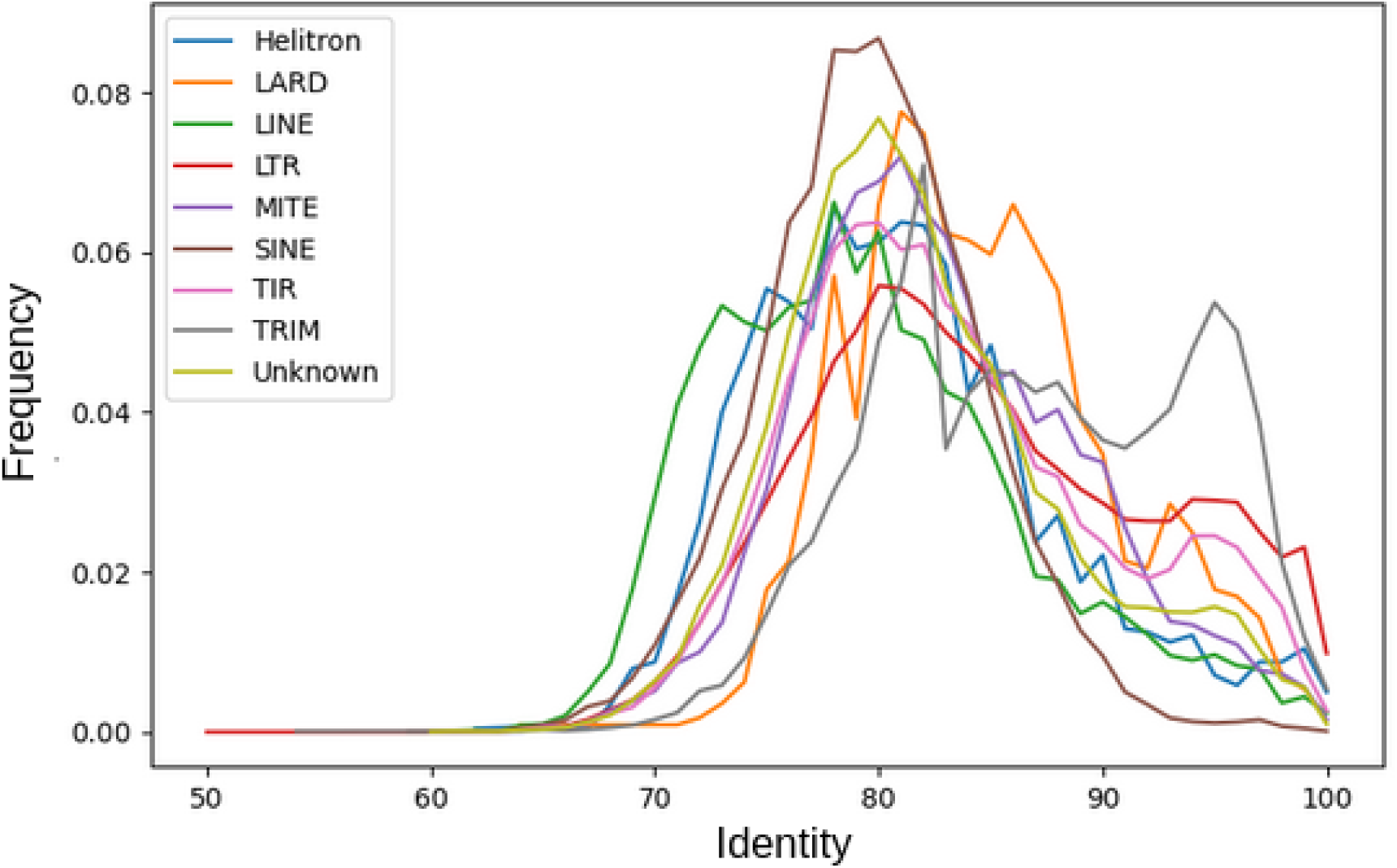
Distribution of the percentage identity of TE copies and their corresponding consensus sequences

TE sequence similarity with their consensus, as well as their insertion positions relative to genes was carried out on a genome-wide scale and further investigated within the environment of the previously defined gene groups.

#### TE insertions in the genome

In apple, among the 441,373 annotated TEs (Daccord et al. 2017), 43,410 (9.8%) were “very recent”; 102,202 (23.2%) TE and were “recent”; and 295,761 (67.0%) were “old”.

In pear, of the 318,639 TEs, 41,027 (12.9%) were “very recent”, 93,676 (29.4%) were “recent” and 183,936 (57.7%) were “old”.

In apple, using the TEGRIP pipeline, we identified 138,957 TEs located within 5kb of genes’ environment (39,977 for pear), regardless of positive or negative DNA strand. Among these TEs, 43,463 (31.3%) were strictly downstream of the genes (9,864 for pear - 24.7%), and 43,608 (31.4%) were strictly upstream of the genes (10,093 for pear - 25.2%). Some TEs were found to overlap with the genes: 5,543 (4.0%) overlapped downstream of the gene (1,866 for pear - 4.6%), and 5,474 (3.9%) overlapped upstream of the gene (1,887 for pear - 4.7%). We identified 33,857 (24.4%) TEs fully inserted into a gene (13,823 for pear - 34.6%) and 7,012 (5.0%) TEs containing a gene (2,444 for pear - 6.1%).

For apple, among the TEs inserted within the 5kb gene environment, 15,794 (11.3%) were tagged “very recent” TEs, 24,699 (17.8%) were tagged “recent” TEs, and 98,464 (70.9%) TEs were tagged “old” TEs. Analysis of TEs within the 5kb environment of genes in the pear genome yielded 41,027 (12.9%) TEs tagged as “very recent”, 93,676 (29.4%) TEs tagged as “recent” and 183,936 (57.7%) TEs tagged as “old”.

#### TE insertions in the gene groups

In apple, within the determined gene groups, non switching over-expressed genes were found to have 2,271 TEs (with 13.5% “very recent”, 17.0% “recent”, and 69.6% “old”) in their environment, while non switching under-expressed were found to have 2,191 TEs (15.3% “very recent”, 19.7% “recent”, and 65.0% “old”) in their environment. Switching genes were found to have 4,584 TEs (11.6% “very recent”, 16.6% “recent”, and 71.8% “old”), and NE 342 TEs (12.0% “very recent”, 18.7% “recent”, and 69.3% “old”). Table 2 reports the number of TE inserted in each gene/TE relationship according the Figure 1 for each differentially expressed gene group. The same informations for pear are in Supplementary data STab3.

**Table 2:** Number of TEs in the different gene/TE relationships (as defined in Figure 1) for the 4 gene groups for the two species. NSO → Non switching over-expressed; NSU → Non switching under-expressed; S → Switching; NE → Not expressed

| Counts | NSO | NSU | S | NE |
| --- | --- | --- | --- | --- |
| apple upstream TE insertions | 719 | 735 | 1513 | 134 |
| apple downstream TE insertions | 710 | 723 | 1498 | 138 |
| apple upstream overlap TE insertions | 82 | 64 | 146 | 12 |
| apple downstream overlap TE insertions | 77 | 78 | 161 | 8 |
| apple subset TE insertion | 683 | 588 | 1262 | 49 |
| apple superset TE insertion | 0 | 3 | 4 | 1 |
| Total apple TE insertions | 2271 | 2191 | 4584 | 342 |
| pear upstream TE insertions | 63 | 65 | 126 | 14 |
| pear downstream TE insertions | 51 | 55 | 114 | 20 |
| pear upstream overlap TE insertions | 8 | 11 | 9 | 1 |
| pear downstream overlap TE insertions | 3 | 7 | 6 | 0 |
| pear subset TE insertion | 110 | 52 | 230 | 17 |
| pear superset TE insertion | 0 | 0 | 0 | 2 |
| Total pear TE insertions | 235 | 190 | 485 | 54 |

Using QuaDS, we investigated a potential over- or under-representation of particular TE in the environment of the 4 gene groups for both apple and pear.

#### Gene groups’ TE enrichment

In apple, non switching over-expressed genes are characterized by an enrichment in TIR (Class II) and SINE (Class I) orders in their environment (Figure 3). They are further characterized by an enrichment of TEs inserted into the genes. Non switching under-expressed genes are characterized by an enrichment in Class I TEs, LTR, LINE and LARD orders, Copia and Gypsy Superfamilies. They are also enriched in “very recent” and “recent” TEs in their environment. Switching genes are characterized by an enrichment in TEs of unknown class and order, as well as by “old” TEs in their environment. Finally, NE are characterized by an enrichment in TEs of LTR order and Copia superfamily in their environment, particularly upstream and downstream of the genes.

**Fig. 3:**
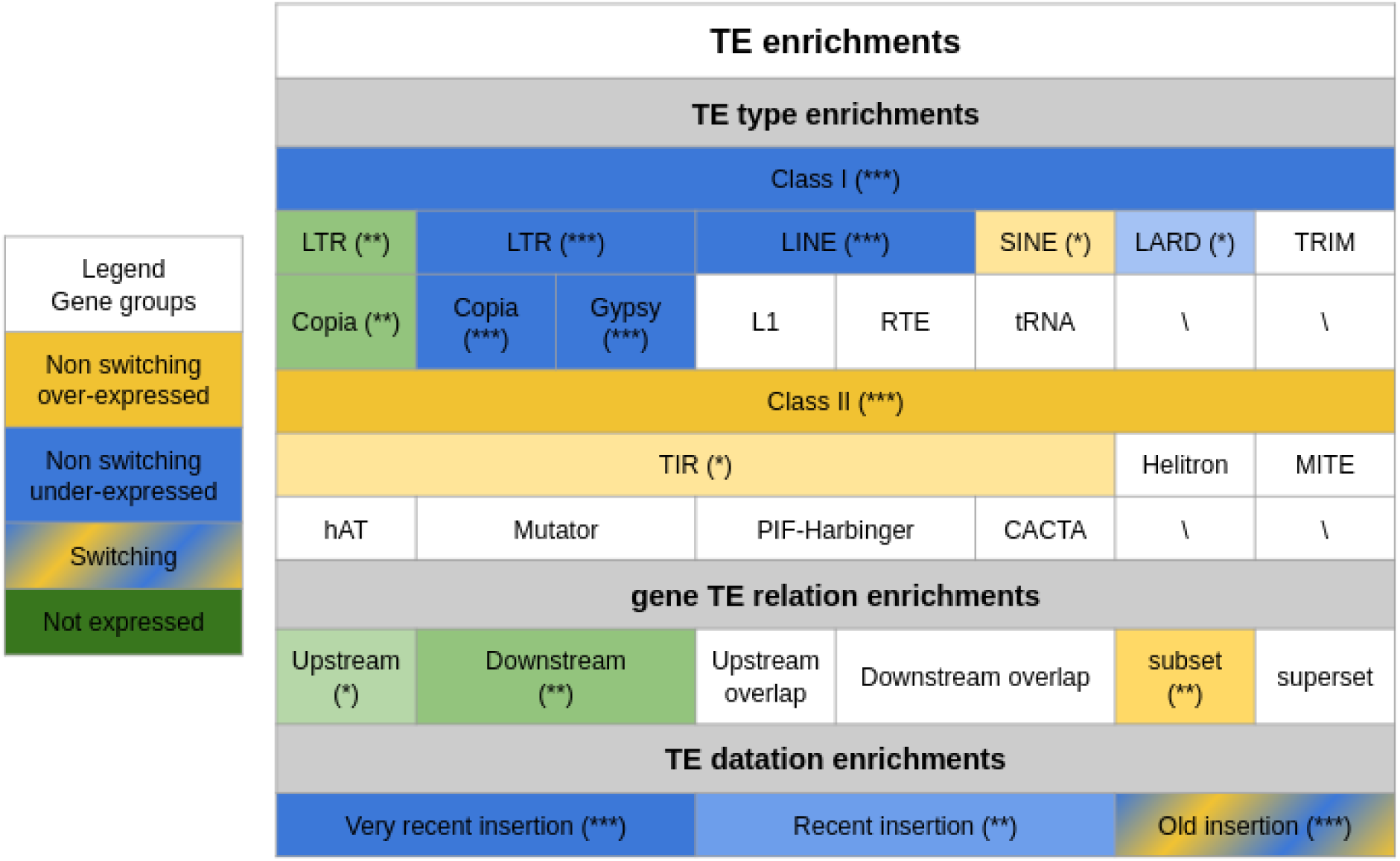
Summary table for the TE enrichment (TE type, gene TE relationship and TE datation) in the different gene groups for apple. Yellow color represents non switching over-expressed over-represented TE Blue color represents non switching under-expressed TE over-represented Combination of yellow and blue colors represent switching over-represented TE Green color represents NE over-represented TE *p*-value are calculated according QuaDS pipeline (*) 0.01 < *p*-value ≤ 0.05 (**) 0.001 < *p*-value ≤ 0.01 (***) *p*-value ≤ 0.001

The enrichment in Copia and Gypsy was found as particularly significant in the non switching under expressed gene group. Copia superfamilly represents only 9.1% of the TEs in the genome and we observed 318 Copia insertions in the 828 non switching gene pairs. These insertions are distributed as follow: 106 insertions in the non switching over-expressed genes environment, representing 33% of the 318 insertions, and 212 insertions in the non switching under-expressed environment, representing 66% of the Copia insertions.

Gypsy superfamilly represents 24.5% of the TEs in the genome. The non switching genes were found to have 432 Gypsy insertions in their environment distributed as follow: 175 insertions in the non switching over-expressed gene environment, corresponding to 40% of the insertions, and 257 insertions in the non switching under-expressed gene environment, representing 60% of the Gypsy insertions.

Analysis of the pear genome yielded similar trends. In this species, non switching under-expressed genes were also found to be enriched in Class I LTR, however, over representation of particular superfamilies could not be established. Within this group, Class I TE LTR located upstream of the gene, upstream overlap and downstream overlap were found to be over-represented. Non switching under-expressed genes were also found to be enriched in Class II Mutator and CACTA superfamilies in their environment. As found in apple, non switching over-expressed genes were also found to be enriched in Class I SINE elements. A summary of the enrichment analysis for pear is presented in table (Supplementary data SFig3).

To identify a potential correlation between the differential expression level within the gene pairs and the number of LTR order TE insertions in the genes’ environment in apple, we classified the genes according to the number of observed TE in their environment.

Our analysis indicated that, in comparison with LTR category 1 genes (logFC=6.70), the logFC of non switching gene pairs increased significantly with the difference in the number of LTR insertions. LTR categories 6, 7, and 9 have more LTR insertions in the environment of under-expressed non switching genes compared with over-expressed non switching genes. LogFC increases significantly by 0.81, 2.08, and 1.89, respectively in these LTR categories (Table 3).

**Table 3:** Table representing the results of the Mixed Model analysis. For each LTR category, the difference in the number of TE between the gene pairs is indicated, as well as the number of TE in the environment of each of the genes of the pair, and the total number of pairs investigated The grey intensities correspond to the difference of LTR TE insertions between the non switching over-expressed and the non switching under-expressed gene groups. The numbers in parentheses in the LogFC Average column correspond to the LogFC average difference between the current category and the LTR category 1. (*) 0.01 < adjusted *p*-value ≤ 0.05 (**) 0.001 < adjusted *p*-value ≤ 0.01 (***) adjusted *p*-value ≤ 0.001

| LTR category | Difference number of TE | Number of LTR in their environment |  | Number of pairs | LogFC Average |
| --- | --- | --- | --- | --- | --- |
|  |  | Non switching over-expressed | Non switching under-expressed |  |  |
| 1 | 0 | 0 | 0 | 283 | 6.70 |
| 2 | 0 | 1 | 1 | 80 | 7.2 (+0.50) |
| 3 | 0 | 2+ | 2+ | 7 | 8.73 (+2.03) |
| 4 | 1 | 1 | 0 | 126 | 6.83 (+0.13) |
| 5 | 1 | 2+ | 1 | 23 | 8.21 (+1.51) |
| 6 | 1 | 0 | 1 | 190 | 7.51 (+0.81)(*) |
| 7 | 1 | 1 | 2+ | 37 | 8.78 (+2.08)(***) |
| 8 | 2+ | 2+ | 0 | 18 | 7.43 (+0.73) |
| 9 | 2+ | 0 | 2+ | 64 | 8.53 (+1.89) (***) |

### 3.6 Methylation profiles

In apple, we investigated a potential link between the expression profiles of our groups of differentially expressed genes and their methylation profiles.

The number of clusters estimated by BPRmeth was found between 6 and 11 depending on the context and the region. The methylation profiles were tagged either hypermethylated, either hypomethylated, or variable.

In what follows, we will first present results obtained for the CG context in the gene sequence region. Next, we will summarize the enrichment of all contexts in all regions.

The different groups of profiles, associated to the different contexts and regions, are presented in Supplementary data SFig4 to SFig8.

We refer also to Supplementary data STab4 to STab6 for the complete QuaDS enrichment analysis. Last, we have to mention that in the pipeline, we used the profile numbers for the QuaDS enrichment, but we summarized the results using the different tags (hypomethylated, hypermethylated or variable) of the methylation profiles (e.g. Figure 4).

**Fig. 4:**
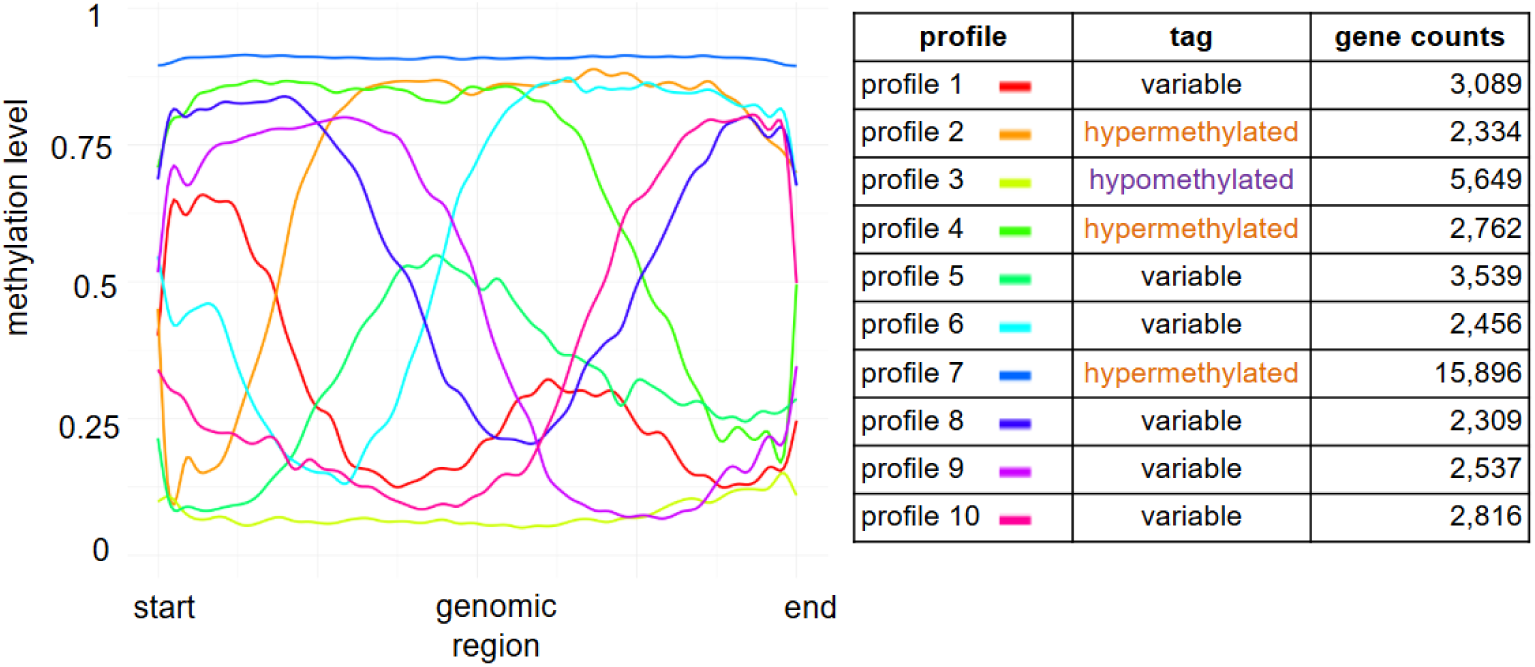
**The 10 methylation profiles**, suggested by BPRmeth, **in the CG context in the gene sequence region**. The 10 profiles have then been tagged either “variable”, “hypermethylated” or “hypomethylated”. Gene counts refer to the number of genes for each profile.

#### 3.6.1 Results for the CG context in the gene sequence region

For the CG methylation context in the gene sequence, BPRmeth suggested 10 methylation profiles (Figure 4).

We then integrated the TE environment of the genes in our analyses (Table 4). The non switching over-expressed genes are enriched in variable methylation profiles and hypermethylated profiles. The variable methylation profile seems to be associated with multiple types of TEs, but the hypermethylated profile is enriched in Class I and LTR. The non switching under-expressed genes are enriched in both hypomethylated and hypermethylated profiles. The hypomethylated profile is statistically associated with Class I, LTR, LINE and unknown orders. When these genes display a Class II, TIR order TE in their environment, they are found enriched in hypomethylated and hypermethylated profiles. Switching genes are enriched in variable methylation profiles. NE genes are enriched in both hypomethylated and hypermethylated profiles. The hypermethylated profile is statistically associated with Class I, LTR, LINE and unknown orders. The hypomethylated profile is correlated with the Class II TE and TIR order.

**Table 4:** QuaDS summary of the over-represented methylation profiles in the gene groups for the CG methylation context in gene sequence region.

| Gene group | Non switching over-expressed | Non switching under-expressed | Switching | Not expressed |
| --- | --- | --- | --- | --- |
| Dataset with all TE | variable + hypermethylated | hypomethylated + hypermethylated | variable | hypomethylated + hypermethylated |
| Dataset with only Class I TE | variable + hypermethylated | hypomethylated | variable | hypermethylated |
| Dataset with only Class II TE | variable | hypomethylated + hypermethylated | variable | hypomethylated |
| Dataset with only unknown class TE | variable | \ | \ | \ |
| Dataset with only LTR order | hypermethylated + variable | hypomethylated | variable | hypermethylated |
| Dataset with only LINE order | variable | hypomethylated | variable | hypermethylated |
| dataset with only TIR order | variable | hypomethylated + hypermethylated | variable | hypomethylated |
| Dataset with only unknown order | variable | hypomethylated | variable | hypermethylated |

#### 3.6.2 General results for all contexts, all regions

The results of all the methylation contexts and regions are presented in Table 5. In contrast to the previous analysis described in Section 3.6.1, we did not filter for TE categories. More detailed results are reported in Supplementary data STab4.

**Table 5:** QuaDS enrichment summary of all methylation contexts (CG, CHG and CHH) in the two genomic regions investigated (500 bp upstream of the gene (ups) and gene sequence (GS)) for all gene groups and all the TE in the genome.

| Gene group | CG ups | CG GS | CHG ups | CHG GS | CHH ups | CHH GS |
| --- | --- | --- | --- | --- | --- | --- |
| Non switching over-expressed | variable + hypomethylated | variable + hypermethylated | hypomethylated | variable + hypomethylated | hypomethylated | variable + hypomethylated |
| Non switching under-expressed | variable | hypomethylated + hypermethylated | hypermethylated | variable + hypomethylated + hypermethylated | \ | hypomethylated |
| Switching | variable | variable | variable | variable + hypomethylated | variable | variable |
| Not expressed | \ | hypomethylated + hypermethylated | hypermethylated | hypomethylated + hypermethylated | \ | hypomethylated |

We found that for the CG context in region 500bp upstream of the gene, non switching over-expressed genes are enriched in hypomethylated and variable methylation profiles, while non switching under-expressed and NE are enriched in variable methylation profiles. For the CHG context, non switching over-expressed are enriched in hypomethylated profiles while non switching under-expressed and NE are enriched in hypermethylated profiles and switching genes are enriched in variable methylation profiles. For the CHH context, non switching over-expressed genes are enriched in hypomethylated profiles while switching genes are enriched in variable methylation profiles.

Results indicate that for the CHG context in the gene sequence region, non switching over-expressed genes are enriched in hypomethylated and variable methylation profiles while non switching under-expressed genes are enriched in hypomethylated, hypermethylated profiles and variable profiles. Furthermore, NE are enriched in hypomethylated and hypermethylated profiles and the switching genes are enriched in hypomethylated and variable methylation profiles. For the CHH context, non switching over-expressed genes are enriched in hypomethylated and variable methylation profiles while non switching under-expressed and NE genes are enriched in hypomethylated profiles and switching genes are enriched in variable methylation profiles.

## 4 Discussion

Following a WGD dated 27 Mya, the apple and pear common ancestor underwent several chromosomal rearrangements and returned to a diploid state, leading to the current 17 chromosomes. This genetic evolution was accompanied by a burst of TEs in apple and pear, with peaks were estimated 21 Mya and 18 Mya for apple and pear, respectively. All these developments impacted the duplicated gene sequences, and affected the preservation of one or both ohnologs within the genomes, or their relative expression. Our results indicate that, based on a survey of the differential expression levels of ohnologous genes in both apple and pear, gene groups displaying particular expression profiles (non switching over-expressed, non switching under-expressed, NE, and switching) can be identified in both species, and show distinct enrichments in GO terms and TE environments. Further investigations in apple indicate that TE types in the genes’ environments may affect the methylation profiles in the gene sequence and within 500bp of the gene start.

Based on our findings, we propose a model describing the evolutionary events that may have led to the systematic over-expression of one gene compared to its ohnolog in the non switching gene group.

In this example, TE insertions may have occurred randomly in the genome throughout the 27 million years post WGD, with a burst of insertions occurring shortly after the chromosome doubling event. TE insertions are often associated with methylation changes around the insertion sites. Our model (Figure 5) presents a TIR order insertion in one ohnologue with, as a potential consequence, a methylation variation in the CG context in the gene sequence, leading to an increased expression of that gene. Sub-data analyses in terms of both class and TE order suggest that insertions of different classes and/or TE orders can induce different methylation profiles both upstream of the gene and within the gene sequence.

**Fig. 5:**
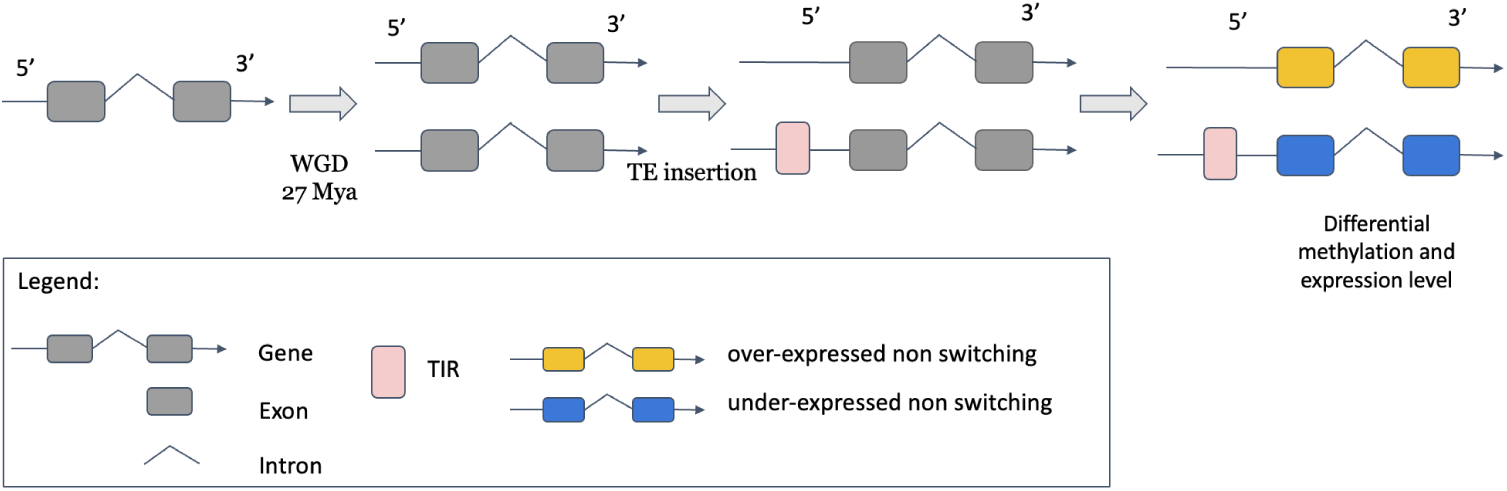
Example of schematic evolution of non switching genes displaying a TIR order insertion in the gene sequence of one ohnolog, with downstream consequences on the methylation profile in the body of the gene in the CG methylation context and eventually on its relative expression level compared to its duplicate. In this example, the non switching under-expressed gene displays a LTR order insertion in the region 500 bp upstream to the gene, resulting in hypomethylation of this region and in the gene sequence in the CG methylation context.

It is important to specify that all analyses performed in this study, besides the differential expression analyses, relate to only one apple genome, the double haploid GDDH13, and one pear species, W65, for the TE annotations and methylation profile. It would be interesting to perform a similar analysis with other diploid apple genomes with their own TE annotation and methylome data in order to validate our approach.

### 4.1 Gene groups are enriched in particular GO terms

In this study we found that non switching genes are enriched in GO terms associated with RNA regulation, metabolism, ribosome biogenesis, and translation in apple, and with mRNA binding in pear. In *Arabidopsis thaliana* and maize, similar GO term enrichments were found for duplicated genes displaying different TE environments (S. Wang and Chen 2019).

In contrast, NE genes in apple are enriched in GO terms associated with plant development, morphogenesis, transcriptional regulation, cell signaling and stress responses, while in pear NE genes are enriched in GO terms associated with enzymatic activities. The various RNA-seq experiments used in our study may not represent all the stages of plant development or all the stresses to which these genes may respond, particularly in pear. In our study, we observed that 0.68% and 0.24% of the duplicated gene were non expressed in apple and pear, respectively. In lotus *Nelumbo nucifera*, 1.15% of the genes duplicated by WGD were found to be non expressed (Shi et al. 2020), while 2.66% duplicated gene pairs were found non expressed in Soybean. (L. Wang et al. 2014). However, no GO enrichment analysis was performed for these two species.

Switching genes in apple are strongly enriched in GO terms associated with transcriptional regulation, gene expression activation, protein phosphorylation and cell signaling, while in pear this gene category is enriched in GO terms associated with transcription factor activity. Their roles in cell development, differentiation, and stress and hormone responses suggest that these genes may be important to environmental adaptation and developmental transitions.

Overall, GO term enrichment analysis highlight a clear functional divergence among gene groups for both investigated species. Non switching genes appear to support essential, housekeeping functions, while switching genes drive adaptive and regulatory transitions in response to developmental and environmental cues.

### 4.2 Variation in ohnologs differential expression is associated with distinct TE environments and methylation profiles

High TE coverage is common in plant genomes. These insertions can impact the expression of genes located near the insertion sites (Hirsch and Springer 2017). As described in Vicient and Casacuberta 2017, TEs can have long-term effects on polyploid genomes. In the apple genome, the gene groups defined in this study and based on the differential expression profiles between ohnologs appear to be characterized partly by different TE environments. Non switching over-expressed genes display more Class II (TIR) in their environment, while non switching under-expressed genes display more Class I (LTR, LINE) in their environment (Figure 5). NE genes are characterized predominantly by the presence of LTR order TE in their environment, while switching genes are enriched in repeated sequences of unknown class. These results are strengthened by the fact that independently, on pear, despite the lower number of genes in each group and the overall lower density of TEs nearby and in the genes, we also find that Class I LTR TE are enriched in the environment of under-expressed genes.

Our results also suggest that, in apple, the differential expression within non switching gene pairs is correlated with the number of TE LTR insertions in the environment of the non switching under-expressed genes. The more TE LTR insertions in the non switching under-expressed environment, the greater the differential expression between non switching ohnologs. This observation indicates that LTR insertions may have a quantitative effect on the reduction of genes expression level. Our results contradict in part the effects of LTR insertions on gene expression in cotton, where LTRs insertions were found to facilitate gene expression while Class II TEs tended to reduce it (Tian et al. 2025).

The association between TEs and the differential expression of syntenic genes had previously been observed in *Malus* (Ze Yu et al. 2024). In this study, Yu and colleagues had showed that differential expression of syntenic genes in the varieties ‘Gala’ and GDDH13 was associated with the presence of TEs proximal to the genes, and suggested that such differential TE insertions could have led to the expression of different phenotypic traits displayed by the two varieties.

In the literature, it has been observed that LTR insertions have an impact on genome size variation and generally tend to induce DNA loss (Ma et al. 2004; Vitte et al. 2007). It was also observed in pear (*P. bretschneideri*) that recent TE insertions (particularly LTR), dated less than 2 Mya, had lead to genome size reduction (Ze Yu et al. 2024). We can hypothesize that this phenomenon may currently be occurring for non switching under-expressed genes and NE genes in both apple and pear.

In fact, LTR insertions in genes’ environment would lead to their pseudogenization (Zhe Yu et al. 2020) following a strong reduction of genes’ expression. Following this hypothesis, we propose that non switching over-expressed genes may become unique singletons in time. Indeed, in apple, within our non switching gene group, we identified 27 pairs of genes in which the under-expressed gene was no longer expressed in the 149 RNAseq experiments, while 7 pairs were observed in pear.

In this study, we investigated particularly the non switching genes group identified from a high number of RNAseq experiments derived from numerous tissues, experimental conditions and varieties. Therefore, we cannot rule out the existence of variety-specific or lineage-specific non switching ohnologues, which differential expression may be induced by particular TE insertions that may have contributed to some of the observed phenotypic differences among apple varieties and species. The identification of such variety or species-specific TE insertions, as observed in oak (Cao et al. 2024), possibly through the development of “panTEome” approaches, would contribute to enhance our knowledge on the effects of TEs on gene expression regulation, and eventually on the emergence of new phenotypes and species.

Gene groups are highly divergent in terms of methylation profiles, both within 500 bp upstream of the genes and within the genes body. Non switching over-expressed genes display mostly variable methylation profiles in the gene sequence and are enriched in hypomethylated profiles 500 bp upstream of the gene for the CHG and CHH methylation contexts. The fact that non switching genes are hypomethylated in their upstream region suggests that non-methylation in the promoter region may favor the gene’s expression. In contrast, non switching under-expressed genes and NE genes are enriched in hypermethylated methylation profiles or display variable methylation profiles 500 bp upstream of the genes’ start in the CG and CHG contexts. The fact that their promoter regions are highly methylated may favor decreased expression (Bird 1995).

We also observed that non switching under-expressed genes displaying hypomethylated profiles in the gene sequence in the CG context were enriched in LTR insertions, which is consistent with the association described in *Arabidopsis thaliana* (X. Zhang et al. 2006). In this study, lower methylation levels in the gene sequence in the CG context correlated with lower gene expression. We therefore hypothesize that the insertion of an LTR in the environment of a gene may result in a decrease of its expression, with additional consequences on the gene sequence methylation profile (compared with other groups of non switching over-expressed, switching, and NE).

It has been observed in soybean and maize that high methylation in the CHG context is inversely correlated with gene expression level (Schmitz et al. 2013; Xu et al. 2019). In our study, non switching under-expressed genes are also enriched in hypermethylated methylation profiles in the CHG context upstream of the gene, as well as in the gene sequence. Similar associations are found for NE genes. We do indeed find a negative correlation between methylation in the CHG context and the expression of duplicated genes, even in duplicated genes containing a LTR in the gene environment.

By analyzing the methylation profile enrichment subdata for differentially expressed gene groups, taking into account TE classes and orders, we found a link between TE environment and methylation profiles. As already observed in rice, maize, and *Brachypodium distachyon*, our study confirms that different TE families can have different impacts on methylation and gene expression profiles (Choi and Purugganan 2018; Noshay et al. 2019; Wyler et al. 2020).

## 5 Conclusion

To conclude, we showed that, in both apple and pear, TE insertions following WGD contributed significantly to their respective genome evolution, and to the currently observed differential expression between ohnologs, possibly through the modification of their immediate genomic environment. We hypothesize that these various TE insertions may have contributed to the emergence of new phenotypic traits, some of which may be associated with plant adaptation, to diversification within the *Malus* and *Pyrus* genus, and may have played a role in speciation.

