## Supplementary material for "LTR transposable elements contribution to the apple genome, methylome and transcriptome evolution in *Malus domestica*": Supplementary_data_captions.pdf

**SFig1** : Barplot of TE coverage by TE order in pear W65 accession.

**SFig2** : Synteny blocks representation (using Circos) of W65 pear accession.

**SFig3** : Summary table for the TE enrichment (TE type, gene TE relationship and TE datation) in the different gene groups for pear.
Yellow color represents non switching over-expressed over-represented TE
Blue color represents non switching under-expressed TE over-represented
Combination of yellow and blue colors represent switching over-represented TE
Green color represents NE over-represented TE
$p$-value are calculated according QuaDS pipeline
(*) $0.01 < p\text{-value} \leq 0.05$
(**) $0.001 < p\text{-value} \leq 0.01$
(***) $p\text{-value} \leq 0.001$

**SFig4** : **The 9 methylation profiles**, suggested by BPRmeth, **in the CG context in the 500 bp upstream region**. The 9 profiles have then been tagged either "variable", "hypermethylated" or "hypomethylated". Gene counts refer to the number of genes for each profile.

**SFig5** : **The 11 methylation profiles**, suggested by BPRmeth, **in the CHG context in the gene sequence region**. The 11 profiles have then been tagged either "variable", "hypermethylated" or "hypomethylated". Gene counts refer to the number of genes for each profile.

**SFig6** : **The 8 methylation profiles**, suggested by BPRmeth, **in the CHG context in the 500 bp upstream region**. The 8 profiles have then been tagged either "variable", "hypermethylated" or "hypomethylated". Gene counts refer to the number of genes for each profile.

**SFig7** : **The 7 methylation profiles**, suggested by BPRmeth, **in the CHH context in the gene sequence region**. The 7 profiles have then been tagged either "variable", "hypermethylated" or "hypomethylated". Gene counts refer to the number of genes for each profile.

**SFig8** : **The 6 methylation profiles**, suggested by BPRmeth, **in the CHH context in the 500 bp upstream region**. The 6 profiles have then been tagged either "variable", "hypermethylated" or "hypomethylated". Gene counts refer to the number of genes for each profile.

**STab1** : Model Mixed for LTR categories.

- **Model Mixed method for LTR categories** : Categories of LTRs based on the number of LTRs in the non switching genes environment and their abundance.

- **Model Mixed output** : Results from the mixed model analysis comparing the mean LogFC values for categories 2 to 9 *versus* the category 1, taking into account the random effect of gene pairs.
- **Model Mixed with p-values ajusted (FDR by BH method)** : details of BH method from the previous Model Mixed output table.

**STab2** : Enrichment tables in GO terms for the three sub-ontologies (BP, MF, CC).

- Sheet 1: Non expressed (NE) vs. switching in apple.
- Sheet 2: NE vs. Non switching overexpressed (NSO) in apple.
- Sheet 3: NE vs. Non switching underexpressed (NSU) in apple.
- Sheet 4: NSO vs. switching in apple.
- Sheet 5: NSU vs. switching in apple.
- Sheet 6: Not differentially expressed (NDE) vs. switching in pear.
- Sheet 7: NSO vs. switching in pear.
- Sheet 8: NSU vs. switching in pear.

**STab3** : TE counts in the environment of different gene groups as a function of TE datation in the W65 pear accession.

**STab4** : Integrative results for profils methylation enrichments of genes groups expression and their environments in all TE for all methylation contexts (CG, CHG, CHH) and all regions (gene sequence or 500 bp upstream).

- Sheet 1: Detailed QUADS output for significative methylation profile overrepresentation (p_value $< 0.05$). The number of methylation profile is in the column 'modalities'. The studied categorical variables are methylation context (column 1), methylation region (column 2) and gene group (column 3).
- Sheet 2: Resumed output of QUADS. The values are the number of the methylation profil enriched in the gene groups, for each regions and methylation context. In yellow are the NSO, in blue the NSU, in green the switching; i, dark green the NE. The color intensity is correlated with significance.
- Sheet 3: Summary of QUADS results (same as Table 5 in main text).

**STab5** : Integrative results for profils methylation enrichments of genes groups expression and their TE environments in class TE (classI or classII ou unclassified) for all methylation contexts (CG, CHG, CHH) and all regions (gene sequence or 500 bp upstream). The caption is the same as STab4.

**STab6** : Integrative results for profils methylation enrichments of genes groups expression and their environments in order TE (LTR, LINE, TIR, Unkmnown) for all methylation contexts (CG, CHG, CHH) and all regions (gene sequence or 500 bp upstream). The caption is the same as STab4.
