## Supplementary figures and images for "LTR transposable elements contribution to the apple genome, methylome and transcriptome evolution in *Malus domestica*"

### Supplementary_data_fig1.png

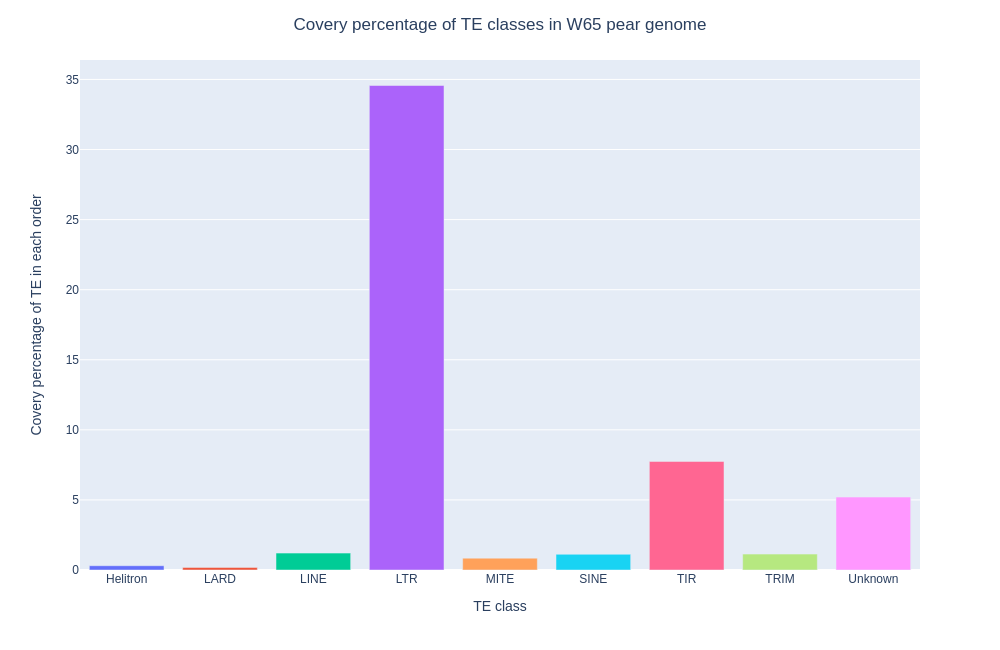

### Supplementary_data_fig2.png

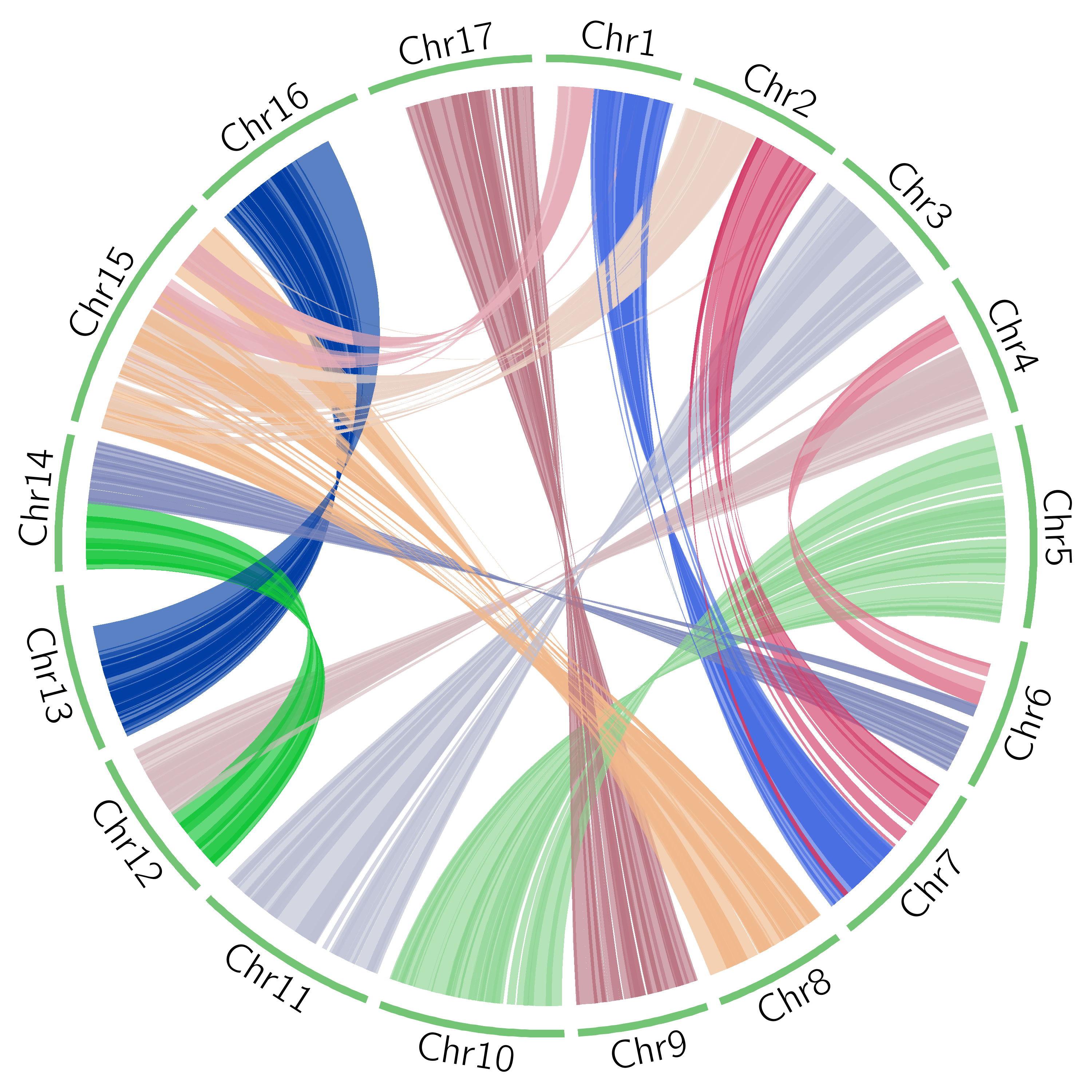

### Supplementary_data_fig3.png

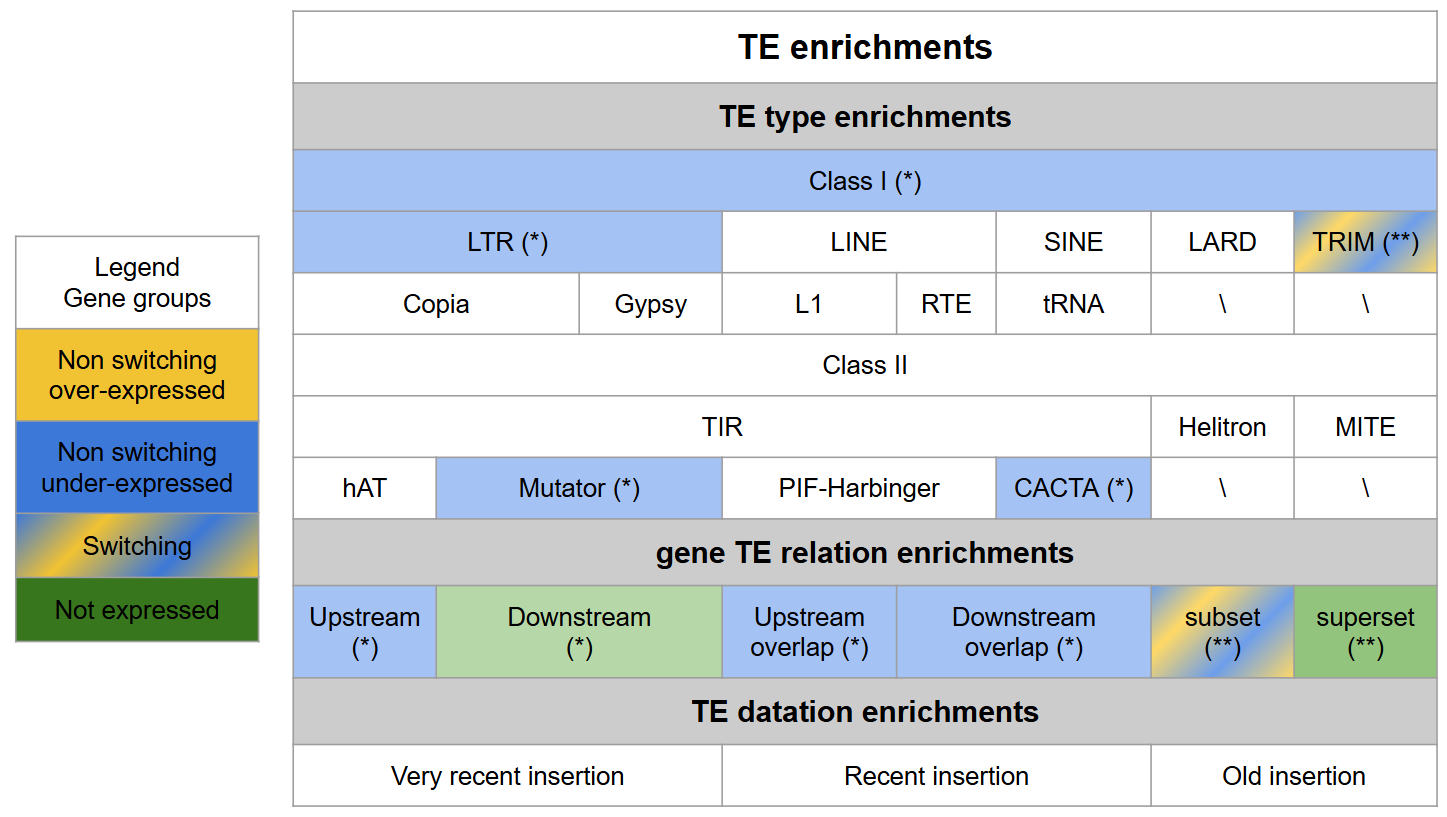

### Supplementary_data_fig4.png

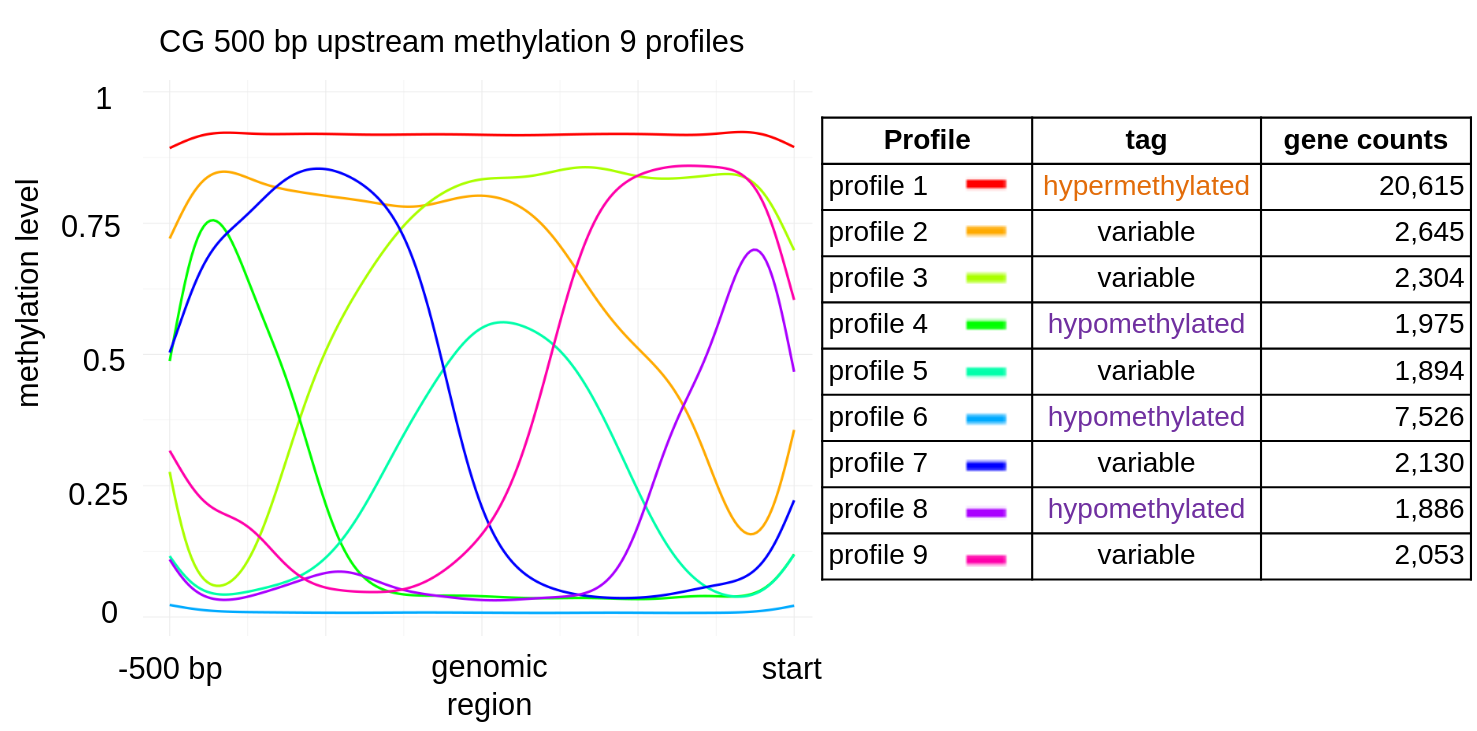

### Supplementary_data_fig5.png

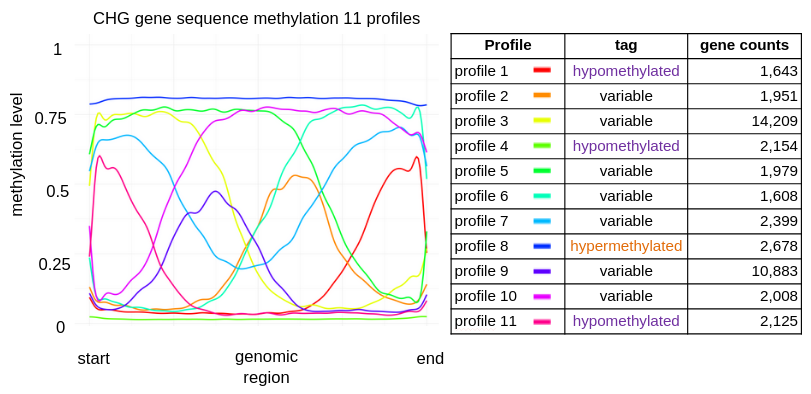

### Supplementary_data_fig6.png

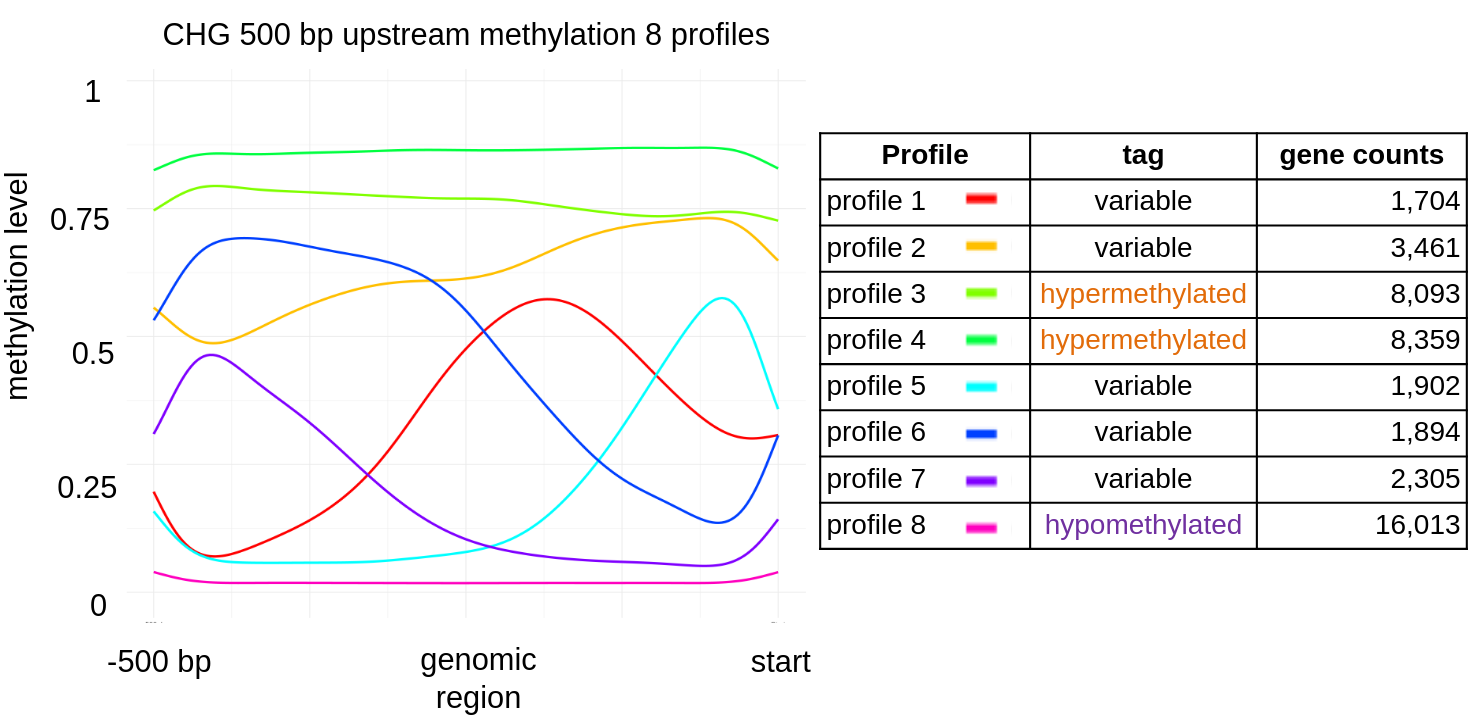

### Supplementary_data_fig7.png

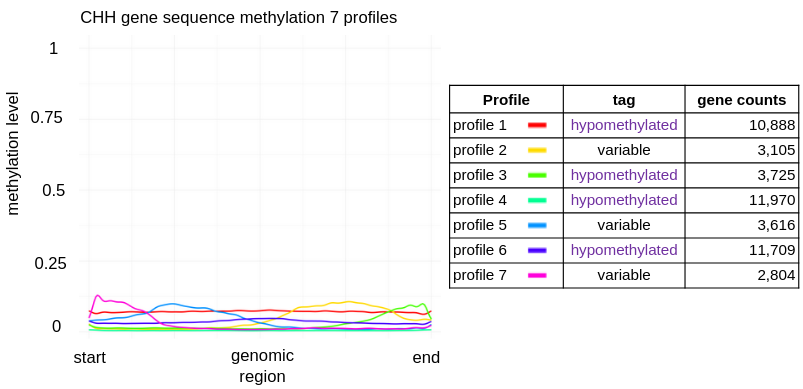

### Supplementary_data_fig8.png

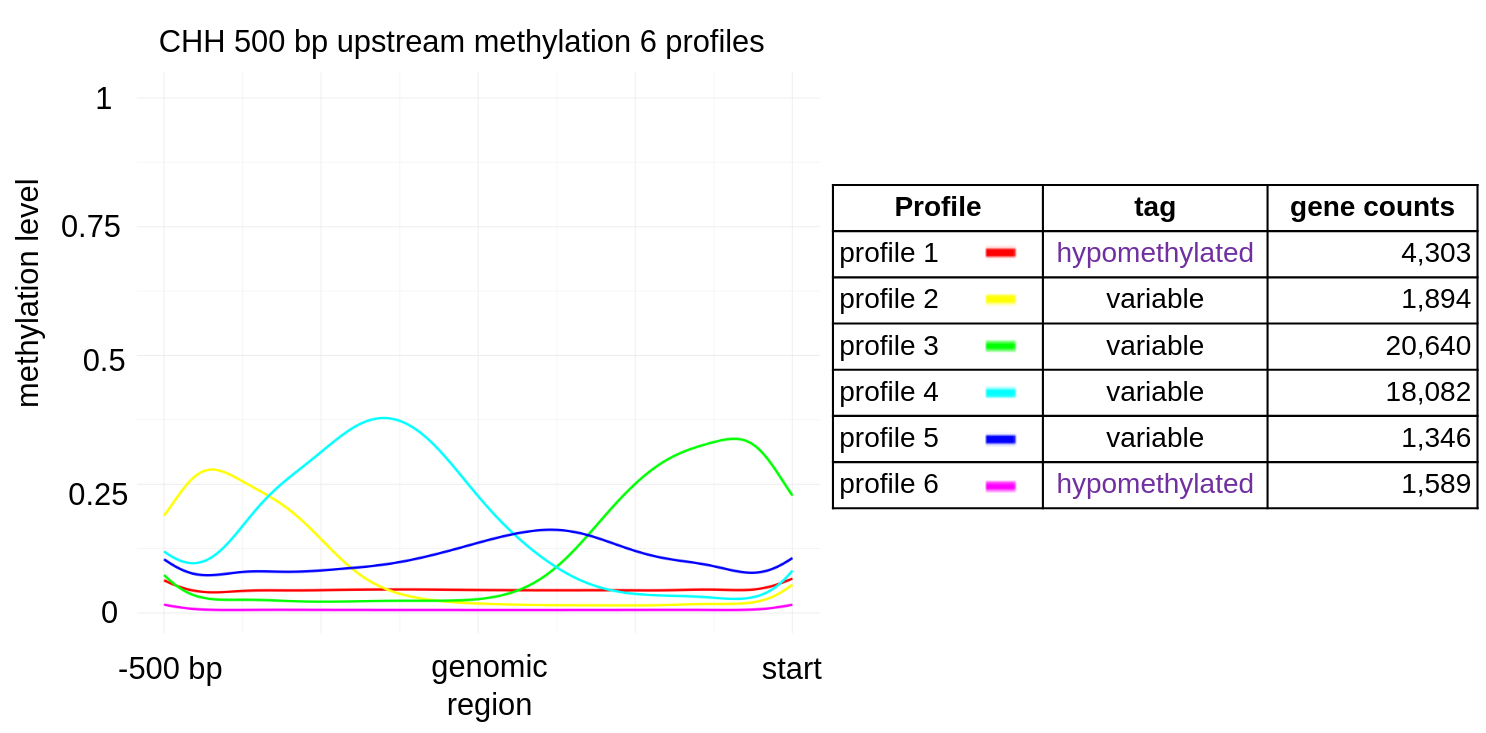
